# Discrimination between *Mycobacterium tuberculosis* and *Mycobacterium bovis* using Fourier Transform Infrared Spectroscopy

**DOI:** 10.1101/2025.01.01.631036

**Authors:** Kevim B. Guterres, Taiana Tainá Silva-Pereira, Rodrigo Oliveira, Carolyn G.J. Moonen, Marcos Bryan Heinemann, Flábio Araújo, Moisés Palaci, Gisele Oliveira de Souza, Nathália Silveira Guimarães, Ana Marcia Sá Guimarães

## Abstract

Zoonotic tuberculosis (TB) caused by *Mycobacterium bovis* (Mb) is a neglected disease that hinders efforts to eradicate human tuberculosis. Developing a rapid, high-throughput diagnostic test to distinguish Mb from *Mycobacterium tuberculosis* (Mtb) isolates could enhance global zoonotic TB diagnostics and surveillance. This study aimed to evaluate the ability of Fourier Transform Infrared Spectroscopy (FT-IRS), using the IR Biotyper® system, to differentiate clinical isolates of Mb and Mtb. Two bacterial inactivation protocols— paraformaldehyde and boiling—were tested using Mtb and BCG strains grown in liquid culture. While both methods allowed FT-IRS analysis, boiling was preferred due to its ease of use and efficiency in biomass recovery. Subsequently, Mtb and Mb isolates were analyzed using FT-IRS, and the resulting spectra were used to construct a sample classifier employing machine learning algorithms. Linear Discriminant Analysis (LDA) and a UPGMA dendrogram demonstrated clear separations between Mtb and Mb ecotypes. Additionally, a classifier built and internally validated using artificial neural networks achieved 99% accuracy in distinguishing Mb and Mtb. Further FT-IRS analysis of few available *Mycobacterium africanum* (Maf) strains demonstrated its capacity to differentiate Maf from Mtb and Mb, expanding its utility in regions where Maf is endemic. This is the first study to apply FT-IRS to distinguish tuberculous mycobacteria. FT-IRS proved to be a highly effective, rapid, and accurate diagnostic tool for differentiating Mb and Mtb strains, with promising applications for other tuberculous mycobacteria such as Maf.

**IMPORTANCE:** Zoonotic tuberculosis (TB) caused by *Mycobacterium bovis* (Mb) remains a major threat due to its clinical similarity to human TB, higher rates of extrapulmonary cases, and resistance to pyrazinamide, complicating treatment. Current diagnostic methods used to differentiate *M. tuberculosis* (Mtb) from Mb and are limited by costs, resource needs, and technical complexity. We developed a method based on Fourier Transform Infrared Spectroscopy (FT-IRS) to differentiate Mb and Mtb clinical isolates with high accuracy. This diagnostic assay offers advantages over traditional molecular techniques by eliminating the need for DNA extraction, requiring less technical expertise, and providing fast, accurate differentiation of Mtb and Mb strains. This innovative approach can improve global diagnostics and surveillance of zoonotic TB.

## INTRODUCTION

The *Mycobacterium tuberculosis* complex (MTBC) comprises 11 ecotypes, each adapted to specific hosts and geographic regions (1). Among these, *M. tuberculosis* (Mtb) is the leading cause of tuberculosis (TB) in humans, accounting for 10.8 million cases and 1.25 million deaths annually (2). Another notable member, *Mycobacterium bovis* (Mb), is the main causative agent of zoonotic TB, with an estimated global burden of 147,000 cases and 12,500 deaths per year (3, 4). Zoonotic TB presents unique challenges to the global control of TB in humans. The disease is clinically undistinguishable from human TB and shows higher proportions of extra-pulmonary cases (5), which complicate diagnosis and delay treatment initiation (4). Additionally, most strains of Mb are intrinsically resistant to pyrazinamide (6), a key first-line drug for TB therapy, subjecting diseased individuals to ineffective treatment in the absence of drug resistance testing. Therefore, to end TB in humans (7), it is crucial to tackle the zoonotic form of the disease.

Major problems in the control of zoonotic TB are its surveillance and diagnostics. The current WHO global TB report does not address zoonotic TB (2) and routine surveillance is absent in 90% of the WHO signatory countries (8). Current diagnostic tests, such as GeneXpert MTB/RIF Ultra and culture, do not differentiate between Mtb and Mb. Culturing Mb is further complicated in Löwenstein-Jensen medium and liquid culture systems, such as BD BACTEC™ MGIT™ 960, due to the absence of pyruvate, which is needed for Mb growth (9). Even if Mb isolates are obtained or new methods are implemented to facilitate their isolation, no rapid tests are available to distinguish Mtb from Mb, which can only be achieved through molecular biology techniques like PCR and whole-genome sequencing (10–12). Therefore, a rapid, high-throughput test is urgently needed to improve individual diagnostics and surveillance of zoonotic TB. This form of the disease is expected to be more prevalent in countries with a high TB burden, where human cases reach hundreds of thousands. In many of these countries, estimates of zoonotic TB are between 2.2% to 8% of all TB cases (13). Therefore, a rapid, high-throughput test would enable more efficient targeted surveillance of zoonotic TB by simplifying the screening of thousands of MTBC isolates.

Infrared spectroscopy (IR) has found wide-ranging applications in microbiology, with Fourier Transform IR (FT-IRS) providing a fast and cost-effective method for distinguishing closely related microorganisms (14). FT-IRS employs a mathematical algorithm to simultaneously capture all wavelengths using an interferometer. This enables rapid and efficient data collection of the molecular signatures of microorganisms (15). This approach has been effectively utilized to differentiate between pathogenic and non-pathogenic strains of *Escherichia coli* (16), to distinguish between methicillin-resistant and - sensitive *Staphylococcus aureus* (17), to detect metabolic changes associated with biofilm formation in *Pseudomonas aeruginosa* (18), to identify *Klebsiella pneumoniae, Legionella pneumophila* and *Acinetobacter baumannii* outbreaks (19–21), among others. Importantly, FT-IRS has been successfully employed to differentiate subspecies of *Mycobacterium abscessus* (22) and to trace the epidemiology of *Mycobacterium chimaera* strains in heater-cooler units used in cardiac surgery (23), attesting its use for the Mycobacterium genus.

FT-IRS has proven invaluable for high-throughput diagnostic applications in clinical settings. Compared to qPCR and WGS, FT-IRS would offer significant advantages in speed, cost-effectiveness, and ease of integration into clinical workflows to distinguish MTBC ecotypes. FT-IRS is less expensive than many molecular biology techniques, making it more accessible in resource-limited settings, and does not require DNA extraction, with sample preparation being fast and easy. In addition, the identification of Mtb and Mb using WGS require skilled personnel for execution and interpretation, which may not always be available. Therefore, the aim of this study was to evaluate the IR Biotyper® system (Bruker Daltonics GmbH & Co. KG, Bremen, Germany) for its ability to differentiate between Mtb and Mb clinical isolates. The developed protocol will enhance the diagnostic accuracy of zoonotic TB, thereby improving the management and control of TB in humans.

## METHODS

### Mycobacteria strains, culture conditions, and ecotype confirmation

A total of 18 MTBC isolates were used in this study, including two laboratory strains (Mtb CDC1551 and Mtb H37Rv), one BCG strain and 15 clinical isolates (Table 1). The bacteria were maintained in Löwenstein-Jensen, Stonebrink or Middlebrook 7H9 (BD, USA) supplemented with 10% OADC [oleic acid (Synth, Brazil), albumin (Invitrogen, USA), dextrose (Sigma-Aldrich, USA), sodium chloride (Sigma-Aldrich, USA), and catalase (Sigma-Aldrich, USA)], 18 mM sodium pyruvate (Dinâmica, Brazil), and 0.05% Tween 80 (Inlab, Brazil) (7H9-OADC-Pyr-Tween) medium. Isolates were stored at -80°C in 7H9-OADC-Pyr-Tween with 20% glycerol.

**Table 1.**
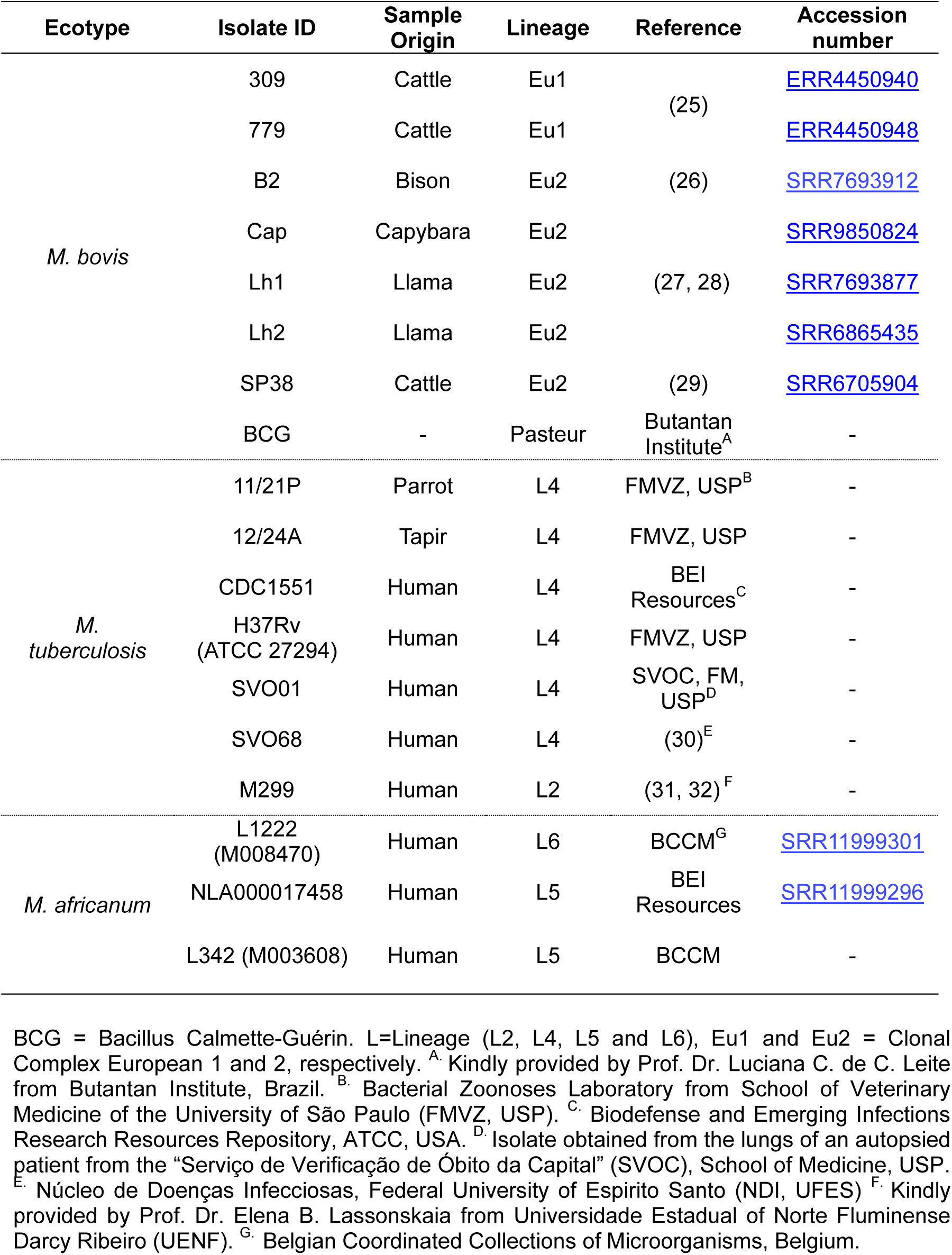
Isolates of the *Mycobacterium tuberculosis* complex included in this study.

To confirm the ecotype of each strain, DNA of the bacteria grown in 7H9-OADC-Pyr-Tween were extracted using a previously described protocol (24) and subjected to a multiplex-PCR to identify RDs (regions of difference) as described (12). RD-based multiplex PCR was also performed before and after experiments to control cross-species contamination. In addition, the genome of all Mb and two *Mycobacterium africanum* (Maf) isolates have been sequenced by our group or others and published previously (Table 1), which further supports their ecotype and clonal complex/lineage identification.

All procedures involving live mycobacteria were performed in a Biosafety Level 3+ Laboratory (BSL3+ Prof. Dr. Klaus Eberhard Stewien), located in the Department of Microbiology, Institute of Biomedical Sciences (ICB), University of São Paulo (USP), Brazil.

### Culture and inactivation protocols

Considering the biosafety requirements to work with tuberculous mycobacteria, an alternative inactivation method to the IR Biotyper kit protocol was sought. For these, a clinical isolate of Mtb (strain SVO01) and a BCG strain were evaluated with different protocols and compared to the kit inactivation protocol using the BCG strain only, which is a BSL2 (biosafety level 2) bacterium. Two protocols were tested: paraformaldehyde (PFA) and boiling. These are commonly used methods of mycobacteria inactivation (33). Briefly, MTBC strains were activated in 7H9-OADC-Pyr-Tween at 37 °C for 5 days in T25 flasks (Greiner Bio-One). Subsequently, 1 mL aliquots were transferred to T75 flasks (Greiner Bio-One) containing 9 mL of 7H9-OADC-Pyr-Tween and cultured for 6-7 days at 37 °C, until an optical density at 600 nm (O.D._600_) of 0.6-0.8. Ten mL of the cultures were centrifuged at 2,000 x g for 15 minutes, resuspended in 1 mL of sterile distilled water, and passed through a 26-gauge needle five times to eliminate bacterial clumps. The suspension was then inactivated using PFA, boiling method or the kit solution (which is 70% ethanol). The PFA protocol consisted in adding the solvent to the suspension to a concentration of 4% and incubating for 40 minutes at 4°C. The suspension was then centrifuged at 2,000 x g for 15 minutes, washed twice with sterile distilled water, and resuspended in a mixture of 50 μL of 70% ethanol and 50 μL of sterile distilled water in an IR Biotyper® suspension vial (Bruker Daltonics GmbH & Co. KG). The boiling method consisted of subjecting the suspension to a dry-block incubation at 95°C for 30 minutes. The suspension was then centrifuged at 2,000 x g for 15 minutes and resuspended in a mixture of 50 μL of 70% ethanol and 50 μL of sterile distilled water in the suspension vial. The suspension containing the BCG strain was subjected to the manufacturer’s inactivation protocol, according to instructions, which consists of ethanol inactivation. Experiments were conducted using 5 replicates of each strain.

Prior to IR Biotyper analysis, aliquots derived after PFA and boiling inactivation were inoculated into rich medium (7H9-OADC-Pyr-Tw and 7H10-OADC-Pyr) and incubated at 37°C for 6 weeks to test the efficacy of inactivation, showing no growth.

### FT-IRS spectra acquisition and data analysis

For the FT-IRS analysis, 24 mL of culture from each strain was used following the same protocol described above. The isolates were analyzed in four to five technical replicates over three independent experiments, whereas the Mtb H37Rv strain was evaluated in six independent experiments. A 15 μL aliquot of each bacterial suspension was spotted onto a 96-well silicon IR Biotyper® plate (Bruker Daltonics GmbH & Co. KG) and dried at 37°C for 15–20 minutes. Additionally, 10 μL each of two IR Test Standards (IRTS1 and IRTS2) were spotted in duplicates.

Spectra were acquired (transmission mode between wave numbers 4000–500 cm^−1^) and processed by OPUS software V.8.2.28 (Bruker Optics, Ettlingen, Germany) on an IR Biotyper^®^ with the corresponding IR Biotyper^®^ software V4.0 (Bruker Daltonics GmbH & Co. KG) for data analysis. After measurement, each spectrum was subjected to a Quality Test (QT). The criteria included for this test were absorption values (range of 0.4-2.0) and signal to noise ratio (R2 <200 and R3 <40). Spectral splicing was performed by default settings to the polysaccharide region of 1300–800 cm^−1^.

Spectral distances were visualized using dimension reduction tools integrated into the IR Biotyper® software. Accordingly, Principal Component Analysis (PCA) or Linear Discriminant Analysis (LDA) with isolate ID set as the target group was used to generate 2D and 3D plots capturing up to 100% of the variance using a maximum of 30 principal components. To visualize differences across all principal components, deviation plots were created, where the solid line represents the mean spectrum, and the shaded area indicates the standard deviation. In addition, a dendrogram using UPGMA algorithm with Euclidean distance was constructed to demonstrates the interrelation proximity of the samples at a certain distance.

### Sample classifier

FT-IRS classifiers enable automated classification of unknown spectra using machine learning algorithms. These classifiers combine a machine learning model, such as an artificial neural network (ANN) or support vector machine (SVM), and an outlier detector (OD). The classifier was developed from labeled spectra, where a set of 242 spectra, representing 7 Mb and 7 Mtb isolates, was used to define two groups. The machine learning model, implemented as a stepwise approach within the software, learns to recognize the specific features of the labeled spectra, classifying unknown samples as either Mtb or Mb based on these patterns. The OD further evaluates how closely the samples match the training set, assigning a reliability score using a “traffic light” system: green for high confidence, yellow for moderate confidence, and red for low reliability, possibly indicating an unknown or significantly different isolate. Accordingly, the dataset of 242 spectra was divided into two groups: a training dataset (total of 124 spectra) consisting of 4 Mtb and 3 Mb isolates and a testing dataset with 3 Mtb and 4 Mb isolates (total of 118 spectra) (Table 2). Of this training dataset, the same machine learning algorithm was used (ANN, 500 cycles) to create a “reduced’’ classifier (of 124 spectra). Next, this reduced classifier was applied to the ‘’unknown’’ testing dataset of 118 spectra resulting in a confusion matrix displaying the accuracy of that reduced classifier.

**Table 2.**
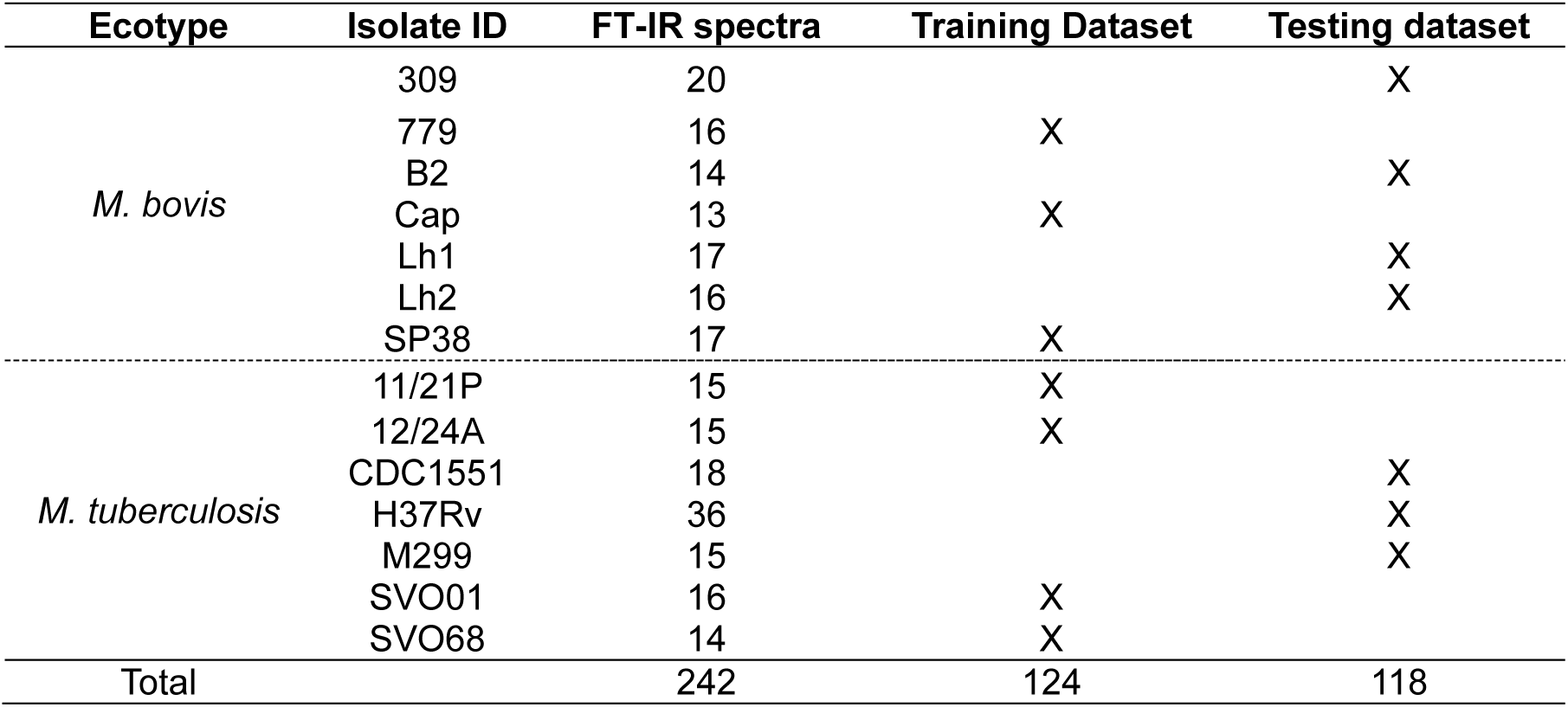
Data of training and testing dataset for the *Mycobacterium tuberculosis* complex classifier.

## RESULTS

### Evaluation of bacterial inactivation protocols

PCA results of Mtb SVO01 and BCG samples subjected to different inactivation protocols are shown in figure 1. Accordingly, Mtb SVO01 and BCG samples cluster separately, but more closely according to the inactivation method used (PFA or boiling) (Figure 1). The PCA graph also shows that the cluster of BCG samples inactivated with the kit protocol appeared in an intermediate position between the samples inactivated with PFA and boiling (Figure 1). Therefore, while different inactivation methods cluster separately, there was evidence of ecotype distinction for both PFA and boiling inactivation protocols.

**Figure 1.**
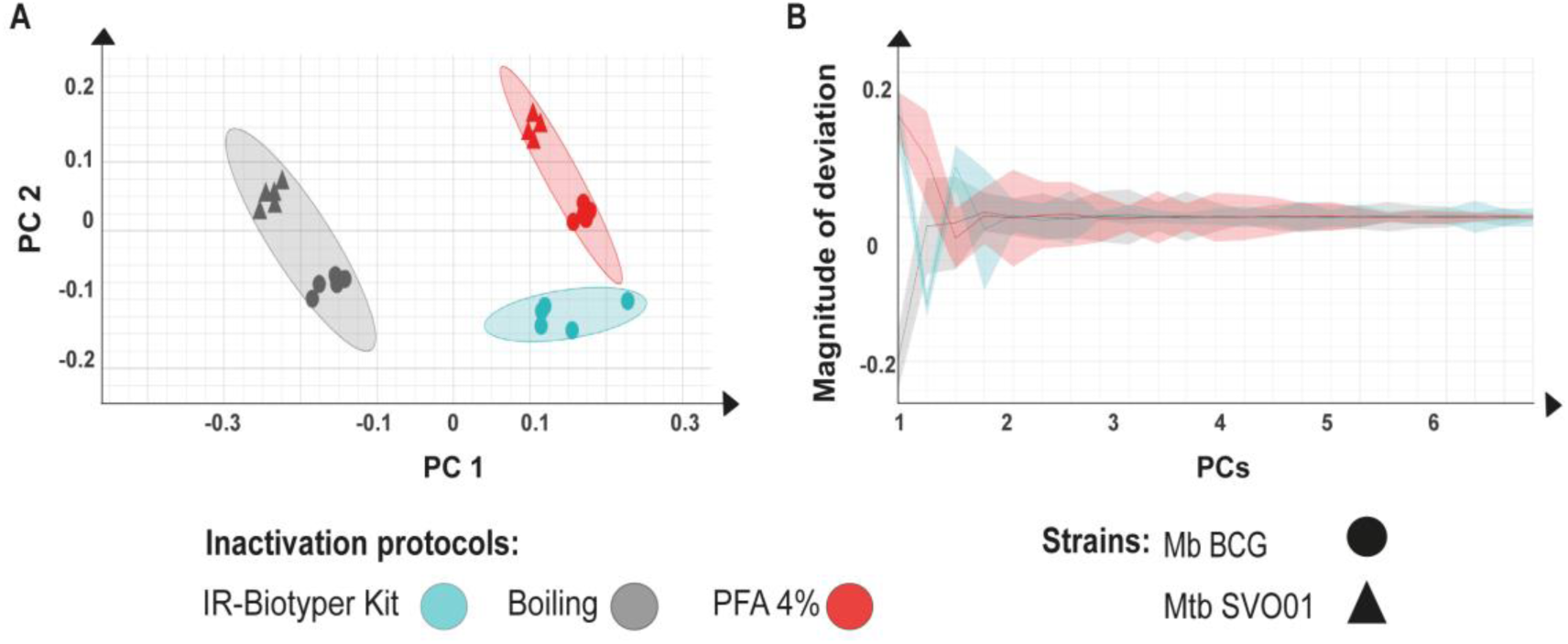
PCA (Principal Component Analysis) and deviation plot of *Mycobacterium tuberculosis* SVO01 and *Mycobacterium bovis* BCG Pasteur inactivated with different protocols and subjected to Fourier Transform Infrared Spectroscopy. **A)** Two dimensions PCA (69.33% of the data variance). Isolates are depicted with different shapes and colors, according to the legend. **B)** Spectra deviation plot for BCG (polysaccharide region of 1300–800 cm^−1^). Solid line displays mean spectra, shade displays the standard deviation.

All samples passed the QT parameters, except for the absorption values, which were below the reference range of 0.4-2 in all samples (mean: 0.2, std: 0.7), except for two samples of BCG inactivated with the boiling method. This suggests that the biomass obtained from 10 mL of cultured media was insufficient to achieve absorption values within the reference range, even though the sample replicates clustered well in the PCA. For this reason, the next experiments were conducted with 25 mL of cultured media and using the boiling method. Boiling was chosen because it separated Mtb from BCG in a similar manner to PFA, is easier to be applied in different laboratories, and requires fewer washes compared to PFA inactivation, resulting in greater amount of final biomass.

### FT-IRS discriminates between Mtb and Mb strains

The LDA of the polysaccharide-based spectra of Mtb and Mb strains shows distinct separation between the ecotypes (Figure 2A and Figure S1). Differences between Mtb and Mb are evident up to LD4, with the greatest distinction seen at LD1 (Figure 2B), which further supports the use of FT-IRS to distinguish the ecotypes. There is a larger standard deviation of Mtb compared to Mb (Figure 2B), likely reflecting the greater distances seen between different Mtb strains compared to Mb strains (Figure 2C and 2D).

**Figure 2.**
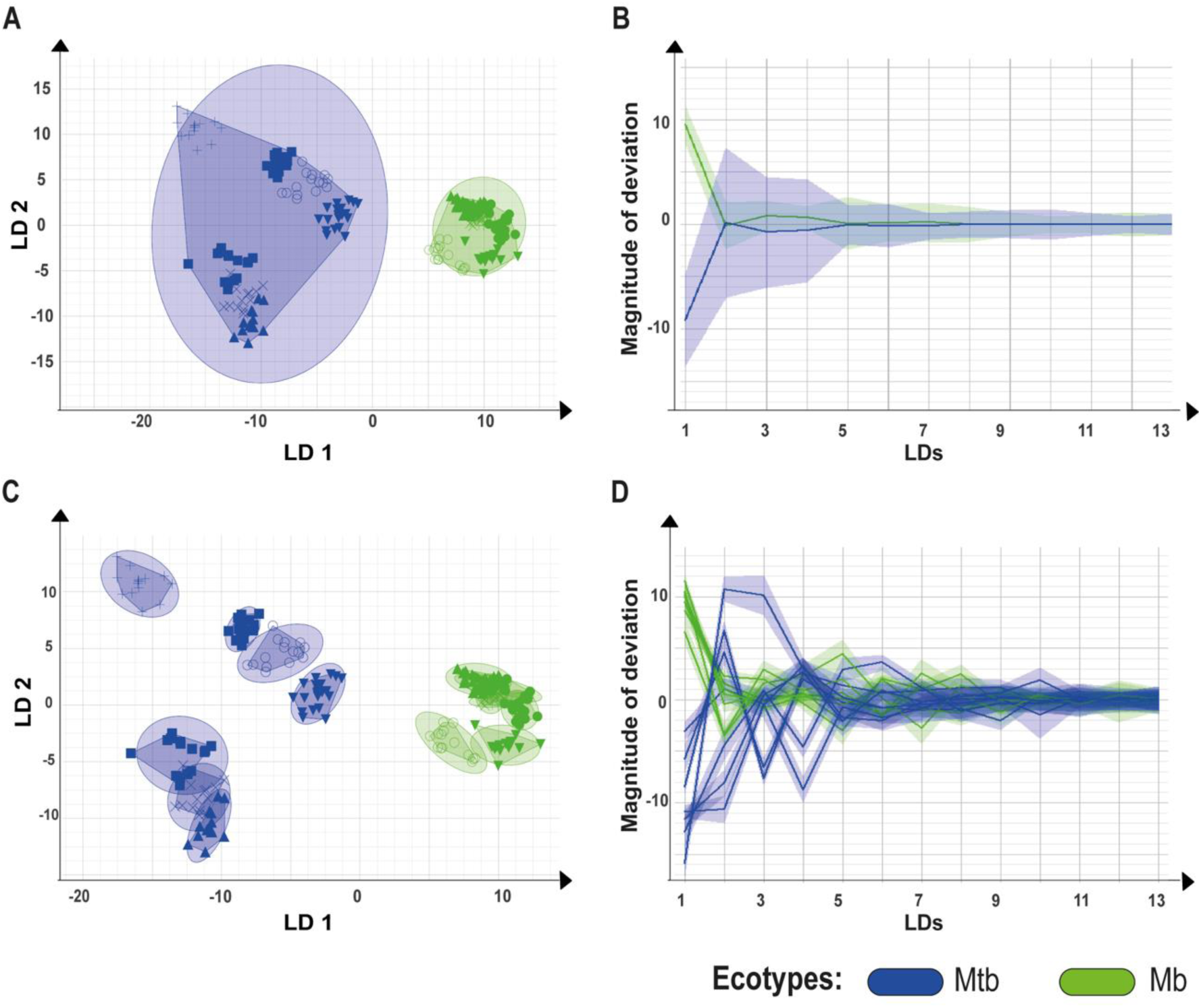
LDA (Linear Discriminant Analysis) and deviation plots of isolates of *Mycobacterium tuberculosis* (Mtb) and *Mycobacterium bovis* (Mb) subjected to Fourier Transform Infrared Spectroscopy. **A)** 2D scatter plot (LDA 30 LDs, 99.1% variance, target group = isolate ID) showing Mtb isolates in blue, Mb isolates in green. Isolates are depicted with different shapes, encircled according to their replicates. X axis is depicting LD1 (59.00%), y-axis is depicting LD2 (15,78%), together displaying 74.78% variance. **B**) Deviation plot displaying Mb in green and Mtb in blue. Solid line displays mean spectra, shade displays the standard deviation. X axis displays different linear components. **C)** 2D scatter plot (LDA 30 LDs, 99.1% variance, target group = isolate ID) showing Isolates depicted in different shades encircled individually by replicates by color, Mtb isolates in blue and Mb isolates in green. X axis is depicting LD1, y-axis is depicting LD2, together displaying 74.48% variance. **D)** Deviation plot displaying Mb individually isolates in green and Mtb individually isolates in blue. Solid line displays mean spectra, shade displays the standard deviation. X axis displays different principal components. Each line represents one spectrum, displaying the variance total of 242 spectra of 14 isolates are displayed.

The dendrogram resulted in a clear distinction between Mtb and Mtb strains into two clades (Figure 3). Within the Mtb clade, there is further separation, represented by the Mtb H37Rv strain, a laboratory strain with unknown number of culture passages, generating what the software calls an incomplete representation of the Mtb group (represented by the yellow color of the branches) given the cutoff value of the dendrogram (Figure 3). While the number of strains from each clonal complex of Mb is low (two Eu1 and four Eu2), we did not observe a clear discrimination between them in the dendrogram (Figure 3).

**Figure 3.**
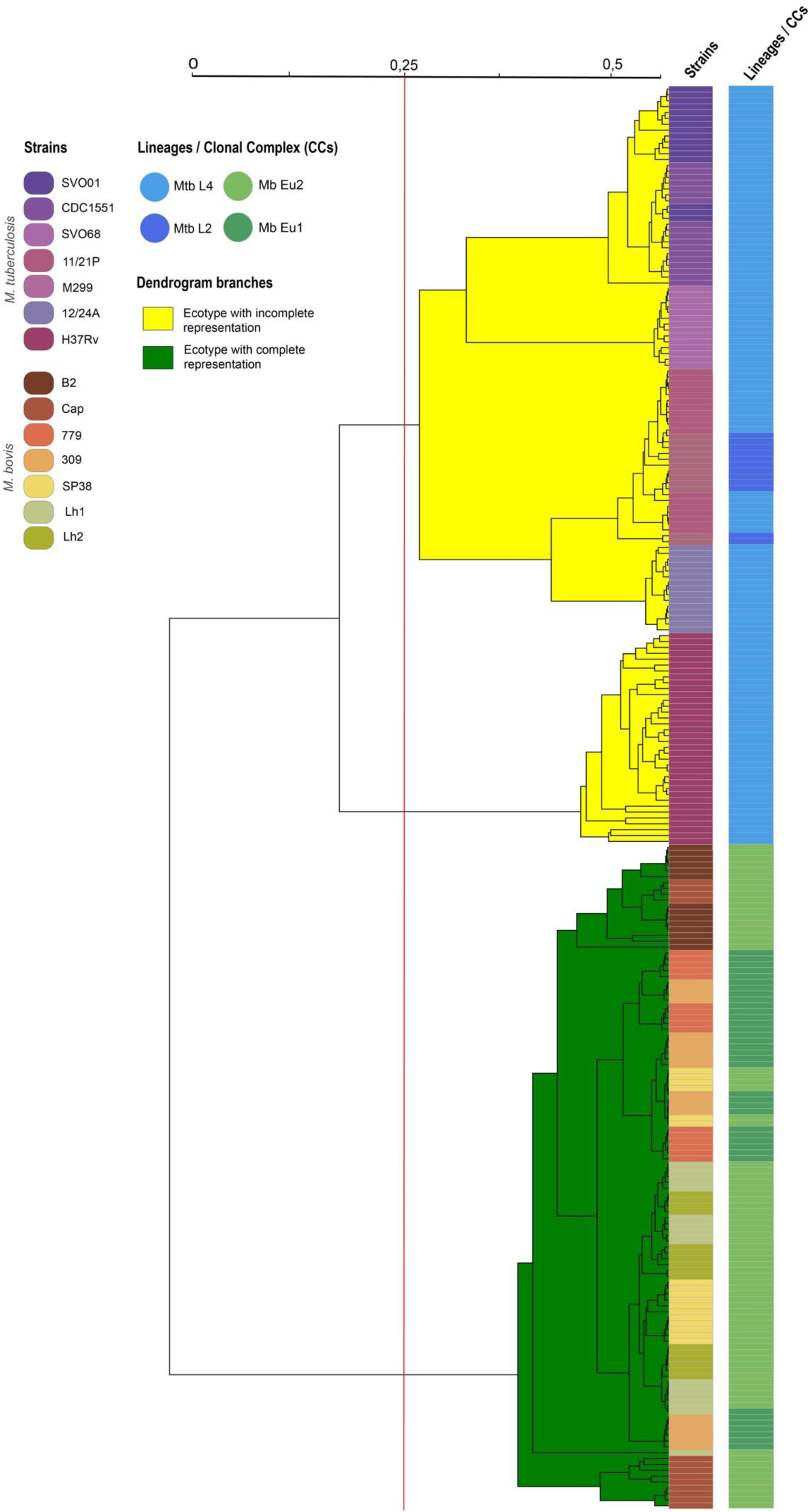
Dendrogram of polysaccharide-based spectra of *Mycobacterium tuberculosis* and *Mycobacterium bovis* isolates subjected to Fourier Transform Infrared Spectroscopy. The wavenumber spectral splicing method was based on polysaccharide range (1300-800 cm^− 1^). Correlation was the exploration method and UPGMA the linkage type. The cutoff value (0.251) of the dendrogram was adjusted based on Simpson’s diversity index × mean coherence, aiding in the visualization of two distinct lusters corresponding to Mtb and Mb strains.

### A classifier for Mtb and Mb

The spectral distance displayed by the Mtb and Mb isolates can be learned by a machine learning algorithm for future automated classification of unknown spectra by the IR Biotyper® software. Accordingly, a classifier was created with the three available machine learning algorithms using a stepwise approach integrated in the IR Biotyper® software. Initially, support Vector Machine (SVM) with a Radial Basis Function (RBF) and SVM with a linear kernel were explored as machine learning algorithms but did not reach >95% accuracy, hence these machine learning algorithms were not further used. A classifier was then created with a total of 242 spectra using the machine learning algorithm Artificial Neural Networks, reaching 100% accuracy (ANN, 500 cycles, Figure 4A). Subsequently, a reduced classifier with a total of 124 spectra was created (100% accuracy, confusion matrix not shown) and validated with other 118 spectra of the testing isolates. Figure 4B shows the confusion matrix of the testing dataset after applying the “reduced’’ classifier, created with 124 spectra. There are two actual classes (labeled isolates) and the output of the predicted classes by the newly created classifier (Figure 4). All spectra, except for one spectrum of isolate Mtb M299, was predicted correctly (Figure 4B). Of the isolate M299, the prediction was 93.3% Mtb, where one spectrum only was predicted as second-best result Mb. Overall, this resulted in the internal validation of this classification model as 99% accurate.

**Figure 4.**
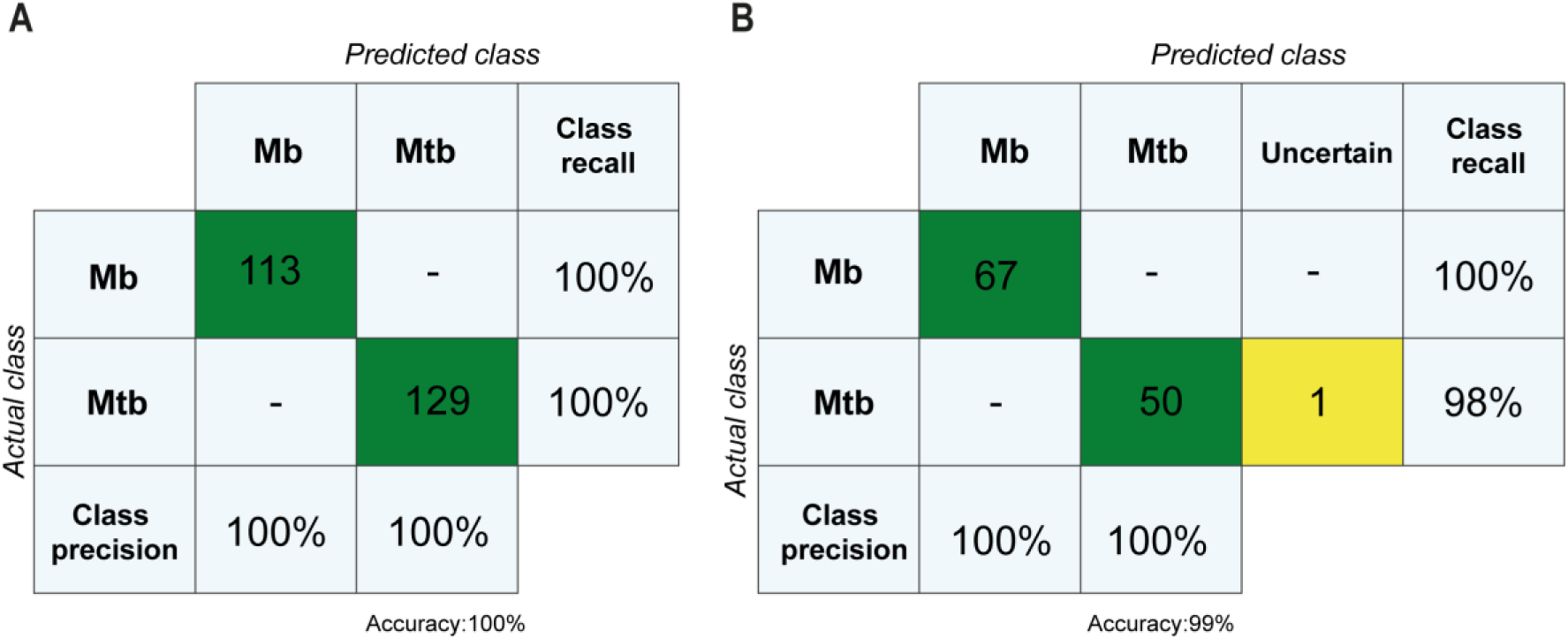
Confusion matrices of classifiers generated with spectra from *Mycobacterium tuberculosis* and *Mycobacterium bovis* isolates subjected to Fourier Transform Infrared Spectroscopy. **A)** Confusion matrix as output from testing the reduced classifier (made with 124 spectra) tested on 118 ‘’unknown’’ spectra in the IR Biotyper® software. The numbers in green depict the number of spectra (67 and 50 for Mb and Mtb, respectively) predicted correctly (matching the actual class), and yellow depict the number of spectra which are uncertain. **B)** Confusion matrix as output from classifier creation of 242 spectra in the IR Biotyper® software. Vertical axis displays the actual class as labels (input given by user), and the horizontal axis displays the predicted classes by the classifier model. The numbers in green depict the number of spectra.

In figure 4A, 100%, (i.e., all 113 spectra) of the Mb (vertical axis, actual class) spectra are predicted (horizontal axis, predicted class) as Mb, resulting in a 100% class recall. The same applied for the Mtb spectra (all 129 spectra), resulting in 100% accuracy of this created classifier. Taken together, these results indicate that the generated classifier created with the total dataset of 242 spectra would be ready to be used for further testing with new isolates as external validation.

### The potential use of FT-IRS to distinguish among additional MTBC ecotypes

Maf is another MTBC ecotype of great importance for human health. However, this ecotype is endemic in West Africa and not in Brazil, where this study was conducted. For this reason, we did not have many isolates of Maf to be evaluated. Only three isolates, acquired through importation from reference databases of microorganisms, were available. Nevertheless, we opted to subject them to FT-IRS analysis and compare them to Mb and Mtb spectra. Figure 5A illustrates the LDA distribution profiles for the three ecotypes using LD1, LD3, and LD5 regions. It reveals three isolated clusters, highlighting the technique’s potential to address the challenge of differentiating other ecotypes. One of the L5 strains exhibited a distinct profile compared to other Maf strains in the initial LD regions (Figure S2), but this difference was mitigated when focusing on a region that shared greater similarity with the ecotype, as observed in LD5 (Figure 5B).

**Figure 5.**
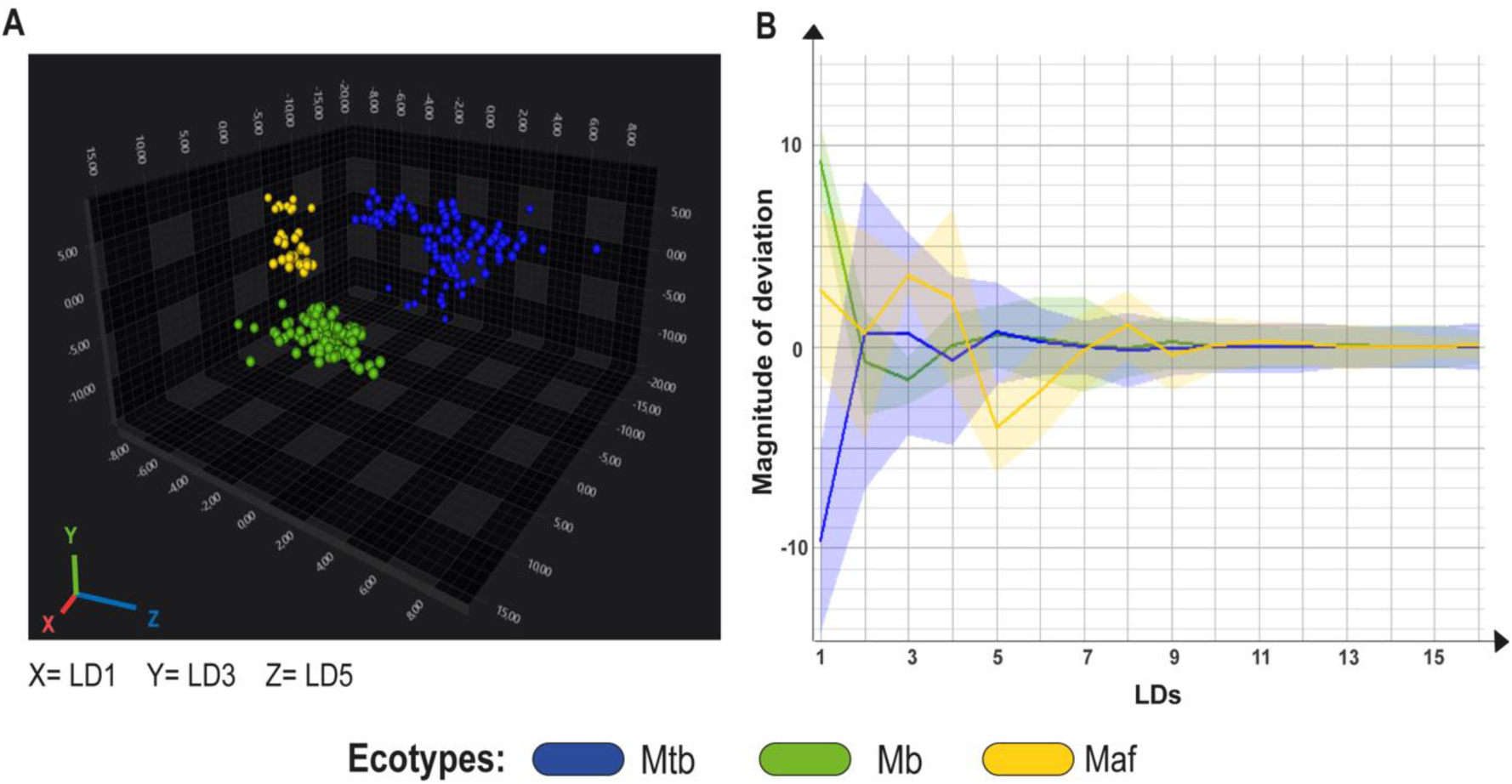
LDA (Linear Discriminant Analysis) and deviation plot of *Mycobacterium tuberculosis* (Mtb), *Mycobacterium bovis* (Mb) and *Mycobacterium africanum* (Maf) isolates subjected to Fourier Transform Infrared Spectroscopy. **A)** 3D scatter plot (LDA 30 LDs, 99.3% variance, target group = isolate ID) showing Mtb isolates in blue, Maf in yellow and Mb in green. X axis is depicting LD1, y-axis is depicting LD3, z-axis is depicting LD5 together displaying 77.99% variance. **B)** Deviation plot displaying Mtb, Maf and Mb isolates. Solid line displays mean spectra, shade displays the standard deviation. X axis displays different principal components. Each line represents one spectrum, displaying the variance total of 255 spectra of 18 isolates are displayed.

## DISCUSSION

This study presents an innovative diagnostic method for differentiating between Mtb and Mb, with potential applicability to other ecotypes within the MTBC. The method offers a rapid, high-throughput test to support diagnostics and surveillance of zoonotic TB and Maf infections. In addition, we showed that the FT-IRS analysis using the IR Biotyper® can be performed on MTBC bacteria cultured in liquid media, which was not the standard protocol (14, 22). The conventional approach required processing samples from solid media. This finding is significant because routine TB diagnostics often rely on automated liquid culture systems, such as the BD BACTEC™ MGIT™ 960, enabling the rapid identification of isolates. Integrating automated liquid media systems with subsequent FT-IRS analysis would enhance uniformity across laboratories, improve quality control, and reduce errors (2, 34).

Due to the alcohol-acid resistance of mycobacteria, a different inactivation method beyond the standard 70% ethanol recommended by the manufacturer was required. Our results demonstrate that the methods tested (PFA and boiling) exhibit distinct clustering behaviors in PCA. This is expected, as boiling induces cell lysis through heat, while PFA likely preserves the cell wall structure by fixing the bacteria (35). For diagnostic purposes, boiling was selected as an efficient and widely applicable method that can be easily implemented in laboratories worldwide. While the default spectral wavelength range of polysaccharide was used herein, it would be valuable to explore other spectral wavelength ranges of mycobacteria inactivated by boiling or PFA. It would be interesting to test if, by maintaining the cells intact with PFA, further differentiation can be achieved among lineages and clonal complexes of each ecotype based on polysaccharide and/or lipids spectra. This is particularly significant because mycobacteria possess a lipid-rich cell wall, with many glycolipids, that are markedly different from those of gram-positive and gram-negative bacteria (36). Thus, this technique holds potential for further applications and exploration.

An interesting result of our study is the larger standard deviation observed in the group spectra of Mtb compared to Mb. Mtb isolates also appeared more dispersed in LDA graphs compared to Mb. These findings suggest that the cell wall of Mtb is more diverse than Mb, with greater variation in polysaccharides (e.g. arabinogalactan and capsular polysaccharides) and/or molecules containing polysaccharides, such as lipoarabinomannan (LAM), trehalose dimycolate (TDM), and phosphatidyl-myo-inositol mannosides (PIM). Interestingly, this greater variability of Mtb spectra aligns with the fact that Mtb strains are more genetically diverse than Mb strains. Despite being clonal and having highly similar genomes, the mean pairwise SNP-distances between Mtb genomes are significantly higher than those of Mb strains (37–39), even when only Mtb L4 strains are considered (40). This is likely because Mb is a more recently evolved member of the MTBC (41, 42). However, since only seven isolates of each ecotype were analyzed, these findings should be further confirmed using a broader panel of Mtb and Mb isolates from diverse global regions.

Mtb H37Rv, the most used reference strain in TB research, was found to cluster separately from other Mtb isolates, suggesting that it may not be an appropriate standard to be used in FT-IRS of the MTBC. Isolated in the 1905 (43), Mtb H37Rv has been distributed to laboratories worldwide and has undergone an unknown number of culture passages. These factors have led to a reduction in virulence (44, 45) and other potentially unknown changes to its phenotype, which may explain our findings.

A sample classifier for distinguishing Mtb and Mb was successfully developed to identify unknown spectra. One spectrum only, from isolate Mtb M299, was misclassified as Mb. However, this isolate would still be assigned as Mtb with over 90% certainty because the other spectra were assigned to the right class. Interestingly, Mtb M299 was the only Mtb L2 used in this work. Due to the higher variability in Mtb spectra, future studies should include more Mtb and Mb isolates to enhance the classifier’s accuracy to 100%.

Maf was included in this study due to its significance as an ecotype infecting humans in West Africa (46, 47). Our results are promising for the potential of FT-IRS in distinguishing this ecotype, demonstrating that this technique can effectively differentiate between Mb and Maf with a level of resolution that even MALDI-TOF (Matrix Assisted Laser Desorption/Ionization – Time of Flight) struggle to achieve (48). This capability underscores FT-IRS’s superior sensitivity to subtle biochemical differences, particularly in the polysaccharide-rich regions of the bacterial cell wall, which seem to be crucial for identifying closely related MTBC ecotypes. Moreover, FT-IRS offers a significant advantage for clinical applications over MALDI-TOF, as it eliminates the need for additional extraction steps (48). Unfortunately, because of the low number of isolates, the sample classifier was not built for Maf.

Limitations of this study include the lower number of strains, especially for Maf, and the lack of other ecotypes and external validation with unknown spectra. Nevertheless, results show that a robust differentiation of Mtb and Mb can be achieved. Furthermore, the created classifier needs external validation. Only an internal validation was done. Ideally, more isolates should be used. However, the created classifier can be easily re-trained with new isolates using the IR Biotyper® software, int the future.

In conclusion, FT-IRS analysis of polysaccharide-based spectra effectively differentiates between Mtb and Mb strains. The sample classifier developed in this study represents a valuable tool for incorporating this technique into routine diagnostics and surveillance of the MTBC. FT-IRS also shows significant potential for distinguishing other ecotypes within the MTBC, potentially offering greater resolution than MALDI-TOF. This advantage likely results from the fact that polysaccharides and lipids provide stronger markers for ecotype differentiation in this clonal group compared to the protein-based approach of MALDI-TOF.

## ACKNOWLEDGMENTS

K.B.G.’s fellowship is funded by São Paulo Research Foundation (FAPESP, 2023/07582-7). T.T.S.P.’s fellowship is funded by the Coordination for the Improvement of Higher Education Personnel (CAPES, 88887.508739/2020-00). A.M.S.G. and M.B.H. are research fellows of the National Council for Scientific and Technological Development (CNPq, 311543/2023-5 and 309146/2017, respectively). This study was funded by FAPESP (2023/13388-9). CAPES (grant #001) provided graduate studies support. The IR Biotyper® kits were provided by Bruker Daltonics GmbH and Co. KG, Germany. We would like to thank the Butantan Institute and Dr. Angela Silva Barbosa for providing the IR Biotyper® equipment.

## AUTHORS’ CONTRIBUTIONS

K.B.G., T.T.S.P., R.O, C.G.J.M., and A.M.S.G. conceptualized and designed the study. K.B.G. performed the experiments. M.B.H., F.A., M.P., G.O.S. and N.S.G. obtained and provided mycobacteria isolates. K.B.G., T.T.S.P., R.O., C.G.J.M. and A.M.S.G. and performed data curation and analysis. K.B.G., R.O., C.G.J.M and A.M.S.G. interpreted the data and drafted the manuscript. Funding acquisition was done by A.M.S.G. All authors edited and reviewed the manuscript.

## SUPPLEMENTARY INFORMATION

**Figure S1.**
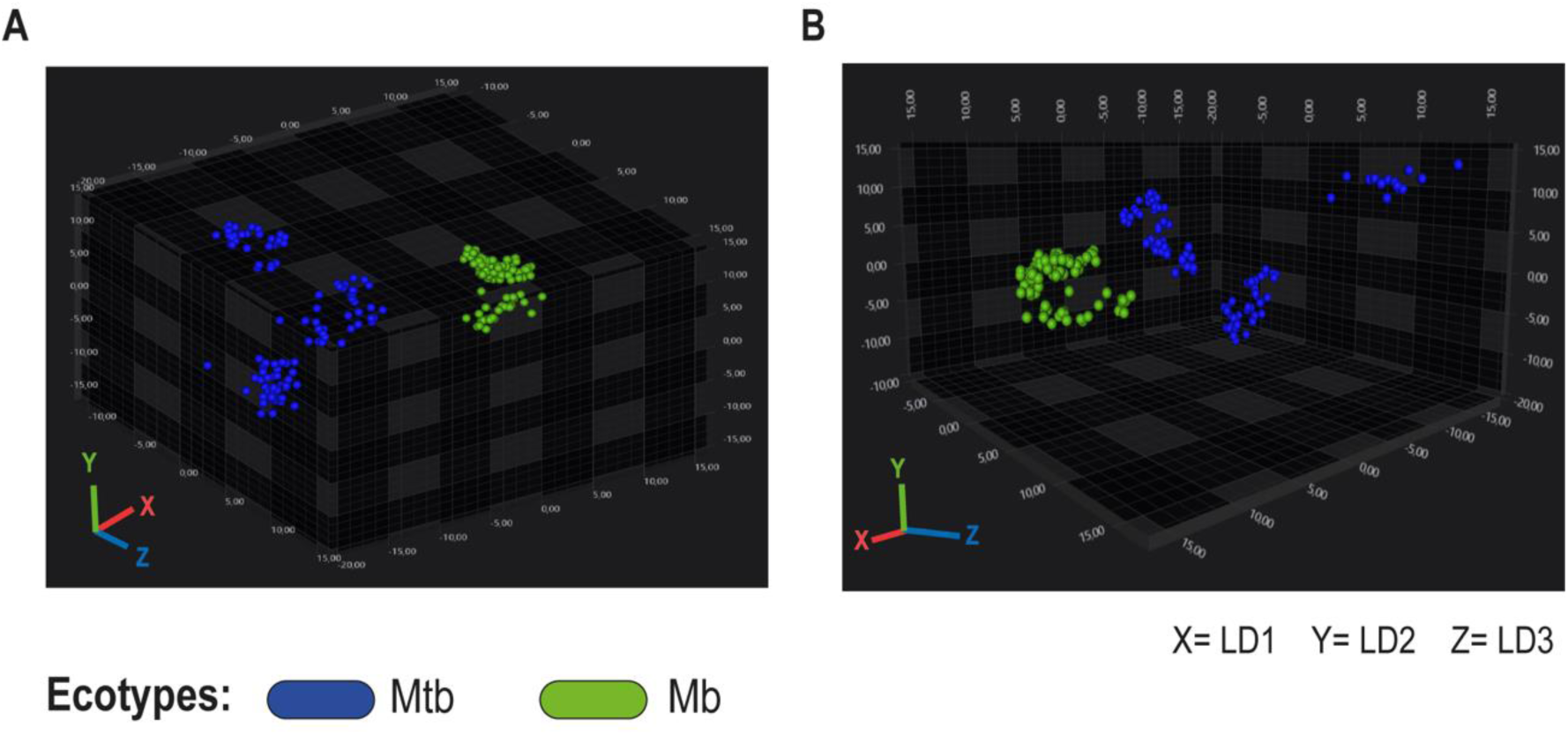
LDA (Linear Discriminant Analysis) of *Mycobacterium tuberculosis* (Mtb) and *Mycobacterium bovis* (Mb) isolates subjected to Fourier Transform Infrared Spectroscopy. 3D scatter plots (LDA 30 LDs, 99.1% variance, target group = isolate ID) showing Mtb isolates in blue, Mb isolates in green. A and B show different perspectives of the same scatter plot. X axis is depicting LD1, Y axis is depicting LD2 and Z axis LD3, together displaying 85.54% variance.

**Figure S2.**
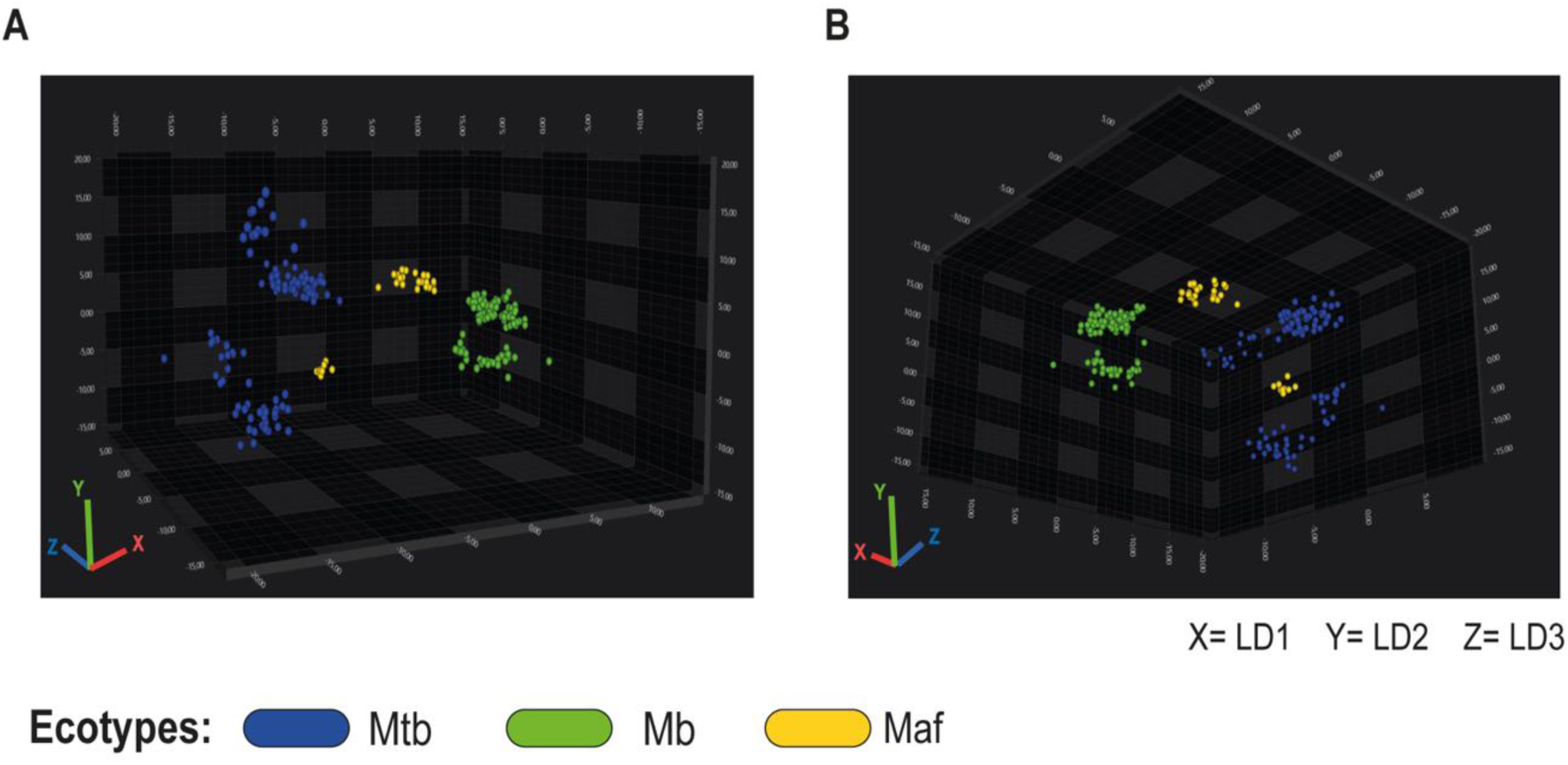
LDA (Linear Discriminant Analysis) of *Mycobacterium tuberculosis* (Mtb), *Mycobacterium bovis* (Mb), and *Mycobacterium africanum* (Maf) isolates subjected to Fourier Transform Infrared Spectroscopy. 3D scatter plots (LDA 30 LDs, 99.3% variance, target group = isolate ID) showing Mtb isolates in blue, Maf in yellow and Mb in green. A and B show different perspectives of the same scatter plot. X axis is depicting LD1, y-axis is depicting LD2, z-axis is depicting LD3 together displaying 82,82% variance

